# Highly accelerated rates of heritable large-scale mutations under prolonged exposure to a metal mixture of copper and nickel

**DOI:** 10.1101/349563

**Authors:** Frédéric J.J. Chain, Jullien M. Flynn, James K. Bull, Melania E. Cristescu

## Abstract

Mutation rate variation has been under intense investigation for decades. Despite these efforts, little is known about the extent to which environmental stressors accelerate mutation rates and influence the genetic load of populations. Moreover, most studies have focused on point mutations rather than large-scale deletions and duplications (copy number variations or “CNVs”). We estimated mutation rates in *Daphnia pulex* exposed to low levels of environmental stressors as well as the effect of selection on *de novo* mutations. We conducted a mutation accumulation (MA) experiment in which selection was minimized, coupled with an experiment in which a population was propagated under competitive conditions in a benign environment. After an average of 103 generations of MA propagation, we sequenced 60 genomes and found significantly accelerated rates of deletions and duplications in MA lines exposed to ecologically relevant concentrations of metals. Whereas control lines had gene deletion and duplication rates comparable to other multicellular eukaryotes (1.8 × 10^−6^ per gene per generation), a mixture of nickel and copper increased rates fourfold. The realized mutation rate under selection was reduced to 0.4x that of control MA lines, providing evidence that CNVs contribute to mutational load. Our CNV breakpoint analysis revealed that nonhomologous recombination associated with regions of DNA fragility is the primary source of CNVs, plausibly linking metal-induced DNA strand breaks with higher CNV rates. Our findings suggest that environmental stress, in particular multiple stressors, can have profound effects on large-scale mutation rates and mutational load of populations.

## Introduction

Germ-line mutations provide the raw material for evolutionary change, but also the genetic variation associated with heritable diseases. Because spontaneous mutations are more often harmful or neutral than beneficial, the accumulation of mutations in the genome has important fitness consequences (Baer et al. 2007; Lynch 2010). The frequency at which mutations are generated, as well as the environmental triggers and selective forces influencing mutation rates are therefore fundamental to biology. Accurately measuring the mutation rate, however, poses a considerable challenge due to the infrequent nature of mutations and the action of natural selection, which eliminates many deleterious mutations to bias the sample of observed mutations. Mutation accumulation (MA) experiments have been particularly effective for directly measuring mutation rates because repeated bottlenecks reduce the effect of selection, allowing all but the most deleterious mutations to accrue over multiple generations (Halligan and Keightley 2009). The comparison of MA experiments with a population experiencing selection can then be used to infer the fitness consequences of new mutations and their contribution to mutational load, ideally by using large populations started with organisms of the same genetic background to eliminate the impact of genotype on mutation rates (Baer et al. 2005; Ness et al. 2015). Studies that conduct whole genome sequencing of MA lines have begun to evaluate the extent to which mutation rates vary across taxa and within species (Schrider et al. 2013; Ness et al. 2015), but few have compared these rates with a population under selection, let alone using the same genetic lineage for this comparison (but see (Flynn et al. 2017)). Furthermore, empirical evidence on the factors underlying mutation rate variation is limited relative to our theoretical understanding (Baer et al. 2007), including the contribution of different environmental conditions, and the long term effects of highly mutagenic environments (Lynch 2016).

The rate of mutations depends on a combination of factors including the amount of DNA damage and the efficacy of the DNA repair machinery, which can both vary under different genetic conditions and environments (Baer et al. 2007; Sharp and Agrawal 2016). DNA damage can be repaired using a multitude of alternative DNA damage response pathways, some of which are more error-prone than others. For example, the two main competing pathways for repairing DNA double strand breaks are homologous recombination (HR) that uses a copy from a homologous template, and a more error-prone nonhomologous recombination (NHR) process called nonhomologous end-joining (NHEJ) that ligates the ends of broken DNA (Ciccia and Elledge 2010; Lam et al. 2010; Carvalho and Lupski 2016). Errors occurring during DNA repair not only contribute to point mutations but are also important sources of copy-number variations (CNVs) – large deletions, duplications and insertions – which can encompass genes and have relevant consequences in cancer and human genetic diseases (Helleday et al. 2014; Sudmant et al. 2015; Carvalho and Lupski 2016). A high propensity for DNA damage or for error-prone repair pathways is therefore likely to elevate mutation rates. Whether these factors are influenced by environmental stressors such as metals to result in higher mutation rates remains largely unknown.

It is established that various exogenous and endogenous stresses induce both DNA breaks and somatic mutations. However, experimental fitness assays have provided indirect and contradictory findings concerning the effects of stress on the accumulation of germ-line mutations (e.g. (Goho and Bell 2000; Joyner-Matos et al. 2011)). Moreover, very few genetic studies have directly investigated the heritable consequences of environmental stresses on the rate of mutations across generations, particularly among multicellular organisms. For example, genetic screens of tandem repeats in either eukaryotic germ-lines or in parents and their offspring have revealed elevated mutation rates upon exposure to air pollution, tobacco smoke, and metals (Somers et al. 2002; Rogstad et al. 2003; Marchetti et al. 2011). Similarly, higher frequencies of CNVs and indels were reported in offspring after parent irradiation (Adewoye et al. 2015). Even scarcer are genomic approaches that utilize MA experiments to assess the variation in mutation rates across environments after multiple generations. MA experiments have revealed that a stressful high temperature increased the rate of short tandem repeats in *Caenorhabditis elegans* measured after 100 generations (Matsuba et al. 2013), and that *Arabidopsis thaliana* grown under salinity stress accumulated about twice as many short insertions and deletions (INDELs) than control lines after only 10 generations (Jiang et al. 2014). However, only one of these past studies surveyed CNVs, which have distinct mutational mechanisms (Lam et al. 2010) that could be more readily induced by stress. It remains unclear whether CNV rates over multiple generations differ across environments and whether they contribute to mutational load.

In this study, we directly estimate genome-wide mutation rates including point mutations, INDELs, and large-scale duplications and deletions under metal stressors in *Daphnia.* This is the first study to estimate large-scale mutation rates under different environmental conditions and under contrasting selection regimes using a single genetic background. Our approach combines two long-term experiments seeded with the same ancestral *Daphnia* lineage: one MA experiment in which selection was minimized and one non-MA population under selection maintained for the entire duration of the MA experiment. This unique design allowed us to directly infer the selective effects on mutations. Additionally, we perform a sequence analysis at mutational breakpoints to inform on the potential source of large-scale mutations and the causes of rate variation across environmental conditions.

## Results

### Mutation accumulation after 100 generations

We sequenced 60 *Daphnia pulex* genomes including 9 MA lines exposed to copper (*Cu*), 9 MA lines exposed to nickel (*Ni*), 9 MA lines exposed to a mixture of nickel and copper (*NiCu*), 24 MA lines maintained in controlled benign conditions (*Con*), and 9 non-MA isolates randomly chosen from a population evolving under selection in benign conditions for the same duration as the MA experiment (Figure 1).

**Figure 1.**
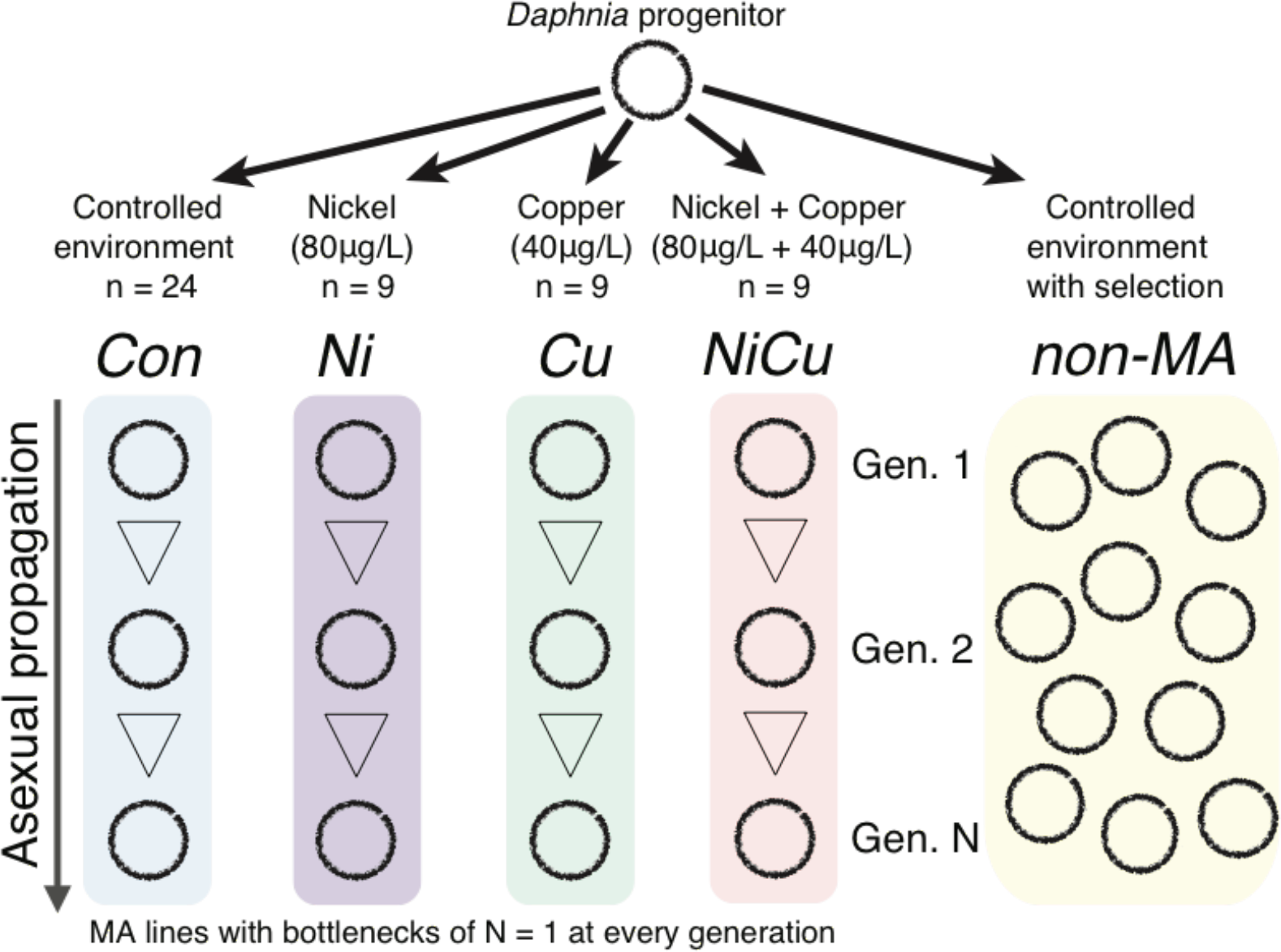
Experimental design. An obligate parthenogenetic *Daphnia pulex* progenitor was used to seed both a mutation accumulation (MA) experiment propagated in four different environments for an average of 103 generations as well as a non-MA population with selection and competition.

The consensus genotype of all MA lines was used to infer the genotype of their common ancestor and the mutations accumulated in each sample. Mutation filtering was calibrated to reduce false positives based on the validation of randomly selected variant calls using PCR and Sanger sequencing (see Methods). After filtering, we detected a total of 916 *de novo* single nucleotide mutations and small (1-50 bp) INDELs, as well as 776 deletions and 406 duplications larger than 500bp (Figure 2; Supplemental Tables S1 and S2). Duplications typically doubled the locus copy number, whereas deletions typically had half the number of reads (Supplemental Figure S1). Genomes with more deletions tended to have more duplications (Pearson’s R = 0.57, p < 0.001), but the number and total length of CNVs per genome were not associated with overall depth of coverage (R^2^ = 0.03, p = 0.09 and R^2^ = 0.02, p = 0.12, respectively). Further, increasing the coverage of two randomly selected MA lines (C01 and C35) did not affect the detection of CNVs.

**Figure 2.**
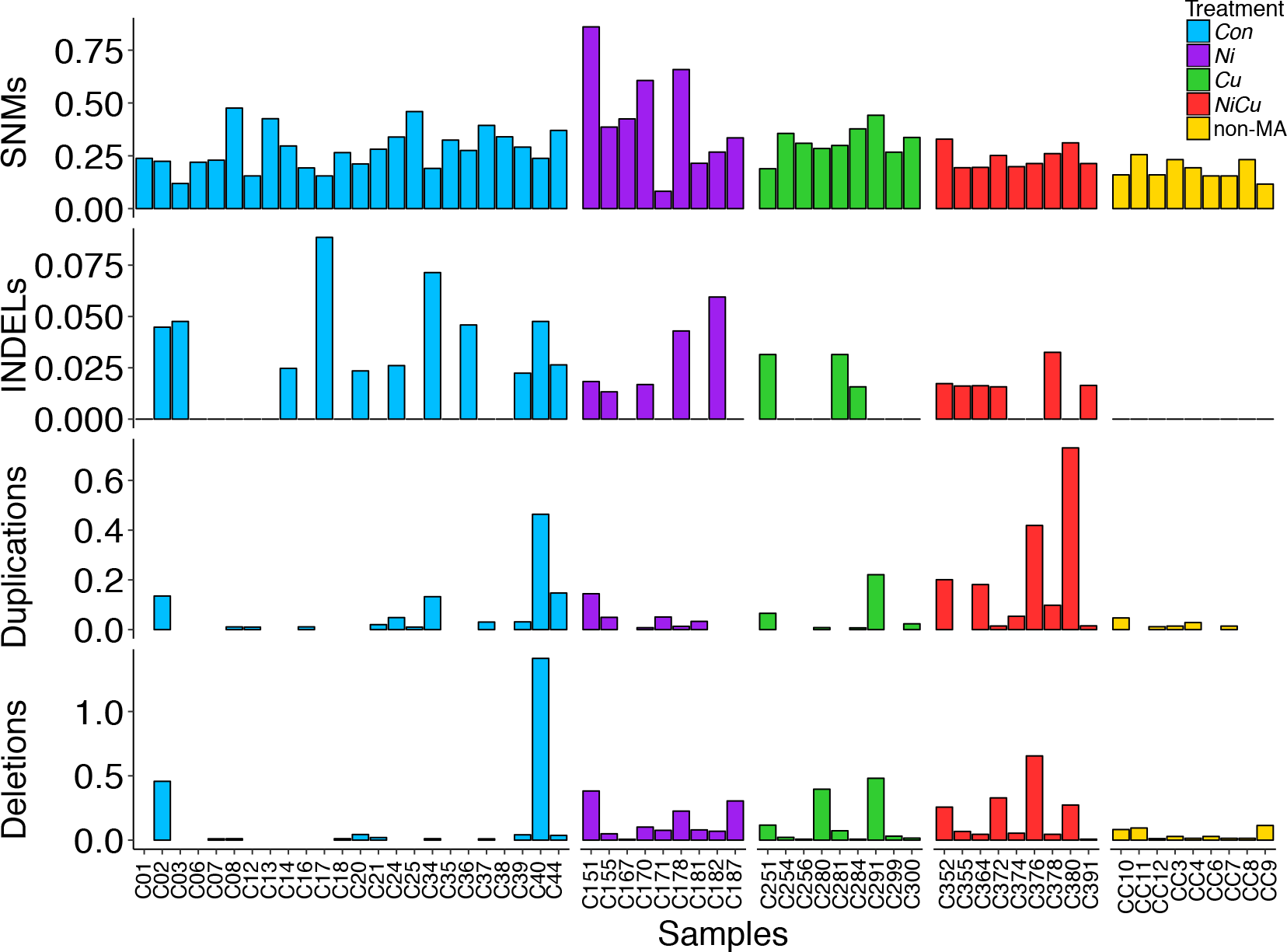
Number of mutations per 100 Mbp per generation detected in each genome. Mutations include single nucleotide mutations (SNMs), small (<50 bp) insertions and deletions (INDELs), and large-scale (>500 bp) duplications and deletions. The number of generations used for non-MA isolates was inferred from a life history experiment.

A total of 243 deletions and 130 duplications overlapped single-copy genes, giving rise to a total of 300 “gene CNVs”, including 180 gene deletions and 139 gene duplications. In addition, there were 177 “partial gene CNVs” that included 136 partially deleted genes and 50 partially duplicated genes (Figure 3; Supplemental Table S3). Many multi-copy genes were found among CNVs, but these were excluded from our gene CNV analysis to limit biases from reads mapping to multiple genomic positions due to the highly duplicated nature of the reference genome (Colbourne et al. 2011; Keith et al. 2016).

**Figure 3.**
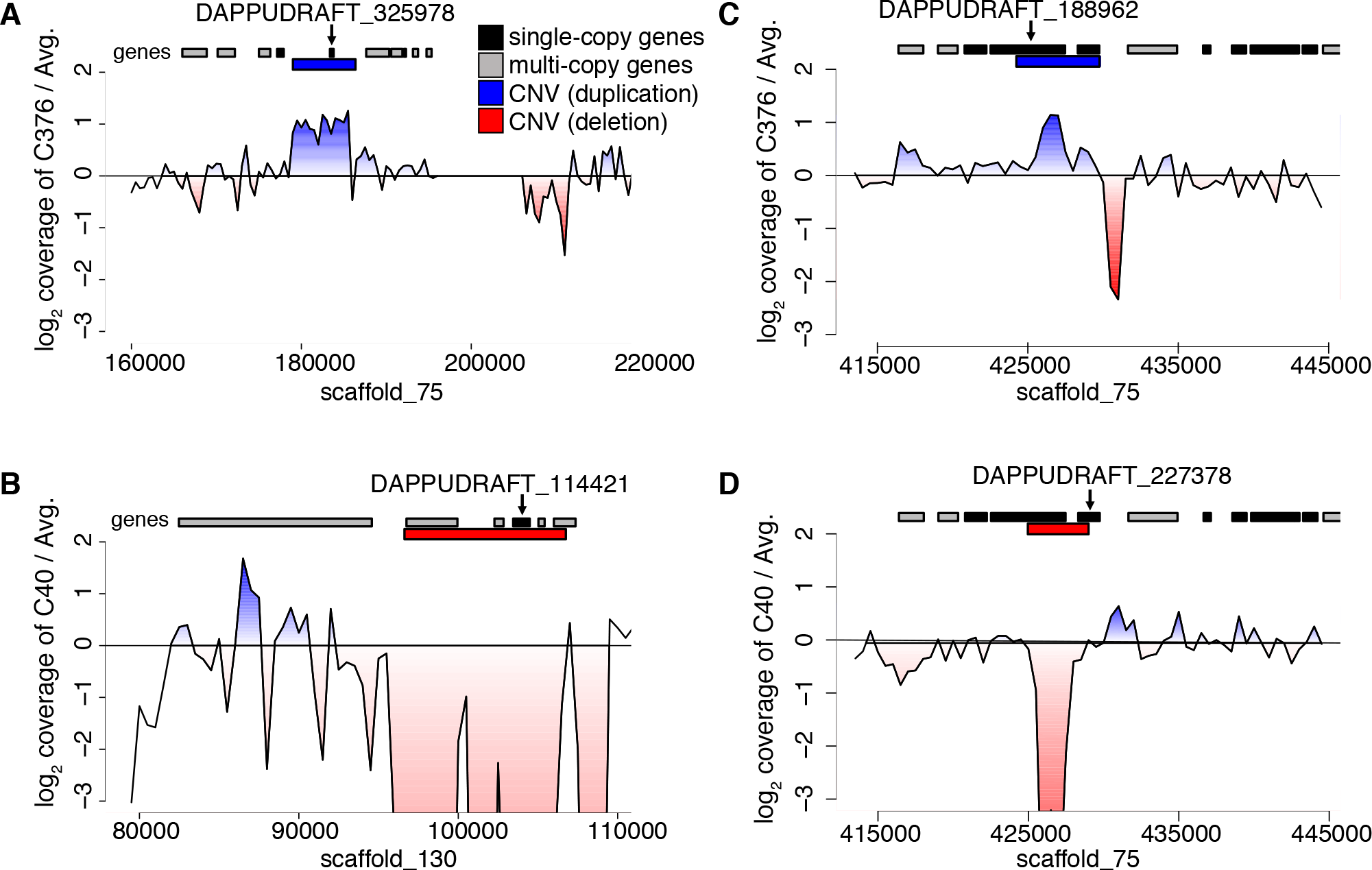
Relative read depth at four CNV loci. Read depth (log_2_ coverage) between genomes with a large-scale mutation (genomic deletions and duplications) and the average of all other MA lines is shown, where a ratio above zero (blue) indicates the focal genome has more coverage than average, and a ratio below zero (red) indicates the focal genome has less coverage. (A) Duplication in the *NiCu* line C376 overlapping an uncharacterized single-copy gene (DAPPUDRAFT_325978). (B) Deletion in *Con* line C40 overlapping an uncharacterized single-copy gene (DAPPUDRAFT_114421) and several multiple-copy genes. (C) A duplication in C376 and (D) a deletion in C40 that lead to a partial gene CNV in the *mre11* gene (DAPPUDRAFT_188962) and a neighboring uncharacterized gene (DAPPUDRAFT_227378).

### Metal stress can increase large-scale mutation rates

A subset of MA lines in our experiment was exposed to metals (copper and nickel) that are prominent environmental stressors in aquatic habitats (Yan et al. 2016). Variation in the mutation rate can arise if cellular stressors due to metals perturb DNA replication, increase DNA damage, or alter DNA repair (Baer et al. 2007). The rates of single nucleotide mutations and INDELs as well as transition/transversion ratios were similar between *Con* lines and the average of all metal-exposed lines (Figure 4).

**Figure 4.**
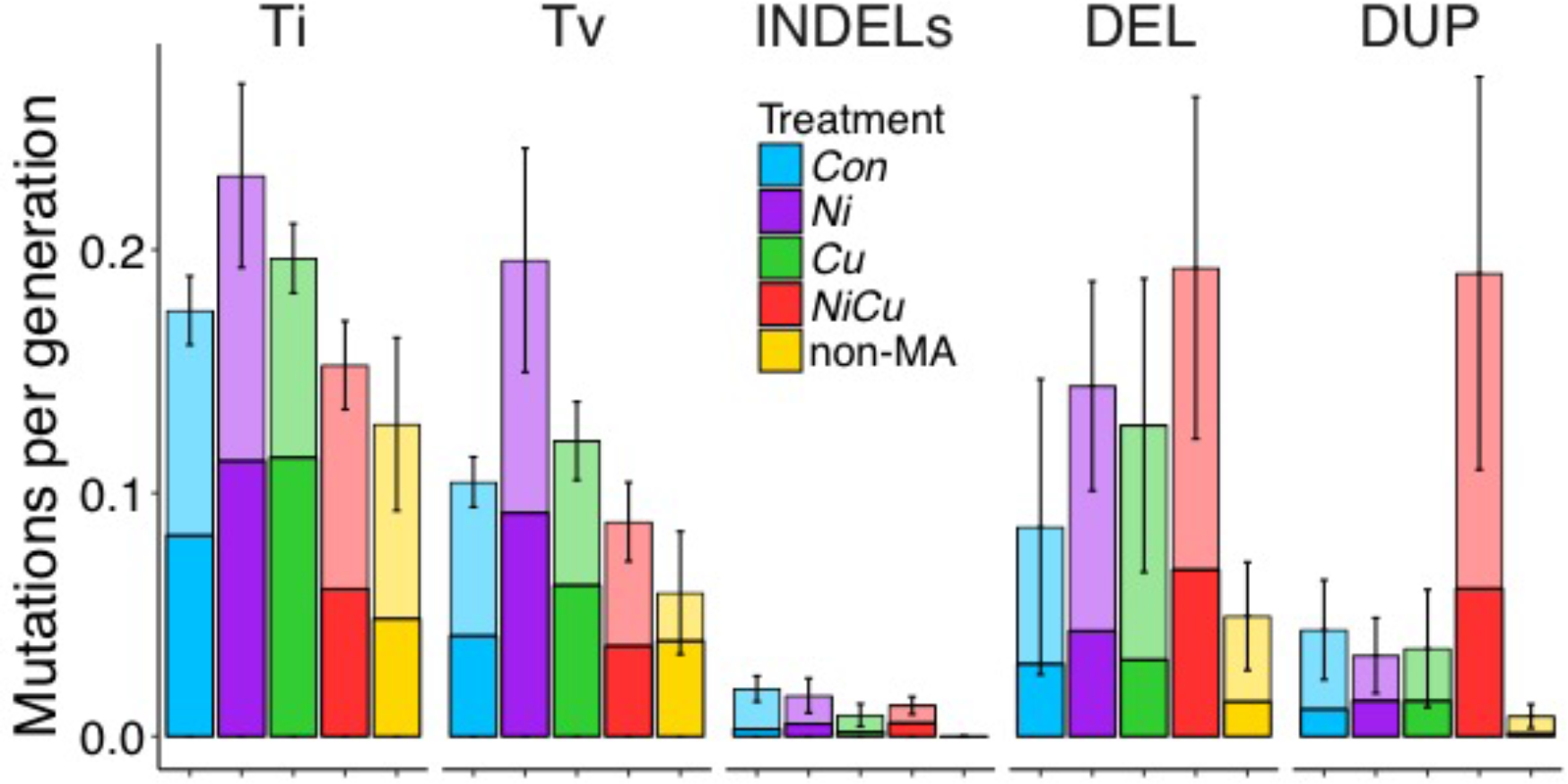
Mean number of mutations per 100 Mbp per generation across treatments and experiments. Stacked bars indicate whether the mutations overlap genes (solid / bottom) or not (transparent / top). Mutations include transitions (Ti), transversions (Tv), small (<50 bp) insertions and deletions (INDELs), and large-scale (>500 bp) deletions (DEL) and duplications (DUP). Standard error bars for each treatment are shown. The number of generations used for non-MA isolates was inferred from a life history experiment.

The highest rate of single nucleotide mutations was observed in lines exposed to nickel, but this was not significantly higher compared to *Con* lines. In contrast, two of the metal-exposed lines (*Ni* and *NiCu*) had significantly greater CNV rates than *Con* lines after Bonferroni correction; the average and standard error (SEM) of CNVs per genome per generation was 0.15 (SEM 0.09) for *Con* lines, whereas *Ni* lines had 1.4x higher rates with an average of 0.20 (SEM 0.06) per genome per generation (Mann-Whitney p = 0.007), and *NiCu* lines experienced over 3.0x higher rates with an average rate of 0.43 (SEM 0.16; Mann-Whitney p = 0.002). *Cu* lines had an average of 0.19 CNVs per genome per generation, but this was not significantly higher than *Con* lines (Mann-Whitney p = 0.069). While deletion rates were significantly greater in each of the metal-exposed lines compared to *Con* lines after Bonferroni correction (all with Mann-Whitney p < 0.01), only the metal mixture *NiCu* had significantly higher duplication rates (Mann-Whitney p < 0.01). Controlling for the number of sites analyzed, the overall rates of CNVs per called site per generation were 6.5 (SEM 4.1) × 10^−10^ for *Con,* 8.9 (SEM 2.8) × 10^−10^ for *Ni,* 8.2 (SEM 4.2) × 10^−10^ for *Cu,* and 19.1 (SEM 0.7) × 10^−10^ for *NiCu.* The elevated CNV rates observed in metal lines remained after accounting for sample size differences across treatments using random permutations (Supplemental Figure S2). CNV rates were not correlated with generation time (Pearson’s R < 0.001, p = 0.99).

### Extensive levels of gene deletions and duplications

To investigate whether the effect of metal stress also extends to functional regions of the genome, we evaluated the impact of CNVs on single-copy genes. As with overall CNV rates, the rate of gene deletions and gene duplications (per gene per generation) varied across samples and treatments (Table 1; Supplemental Table S1).

**Table 1:** Mean CNV, deletion (DEL) and duplication (DUP) rates ± standard errors for the whole genome and for single-copy genes per generation are shown for each treatment. Rates include all genome-wide CNVs per generation, all gene CNVs including partially deleted and duplicated genes, and complete gene CNVs excluding partially duplicated and deleted genes. Rates for the non-MA isolates were calculated using conservative estimates of generations derived from a life history experiment.

| | Genome-wide ($\times 10^{-1}$) | | | Genes ($\times 10^{-6}$) | | | Complete Genes ($\times 10^{-6}$) | | |
| --- | --- | --- | --- | --- | --- | --- | --- | --- | --- |
|  | CNVs | DEL | DUP | CNVs | DEL | DUP | CNVs | DEL | DUP |
| *Con* | 1.5±0.9 | 1.0±0.7 | 0.5±0.2 | 3.1±1.7 | 2.1±1.4 | 0.9±0.4 | 1.8±1.0 | 1.0±0.7 | 0.8±0.3 |
| *Ni* | 2.0±0.6 | 1.6±0.5 | 0.4±0.2 | 5.9±2.8 | 4.0±2.1 | 1.8±1.4 | 3.9±2.0 | 2.5±1.5 | 1.5±1.1 |
| *Cu* | 1.9±1.0 | 1.4±0.7 | 0.4±0.3 | 3.8±1.9 | 2.7±1.3 | 1.1±0.7 | 2.5±1.3 | 1.9±1.0 | 0.6±0.3 |
| *NiCu* | 4.3±1.6 | 2.2±0.8 | 2.2±1.0 | 12.4±3.6 | 6.4±2.4 | 6.0±2.6 | 7.8±2.5 | 3.3±1.4 | 4.5±1.9 |
| non-MA | 0.6±0.2 | 0.4±0.2 | 0.1±0.1 | 1.4±1.2 | 1.1±1.3 | 0.3±0.2 | 1.3±1.3 | 1.1±1.3 | 0.1±0.2 |

The mean deletion and duplication rates overlapping genes (both partially and completely) in *Con* lines were 2.1 (SEM 1.4) × 10^−6^ and 0.9 (SEM 0.4) × 10^−6^, respectively. The combined rate amounts to 3.1 (SEM 1.7) × 10^−6^ CNVs per gene per generation. *Ni* lines had higher gene deletion (4.0 (SEM 2.1) × 10^−6^) and gene duplication (1.8 (SEM 1.4) × 10^−6^) rates, but these were not significant after Bonferonni correction. Similar results were found for *Cu* lines with a gene deletion rate of 2.7 (SEM 1.3) × 10^−6^ and a gene duplication rate of 1.1 (SEM 0.7) × 10^−6^. In contrast, *NiCu* lines had gene CNV rates four times higher than *Con* lines at 12.4 × 10^−6^ (Mann-Whitney p < 0.001), with a gene deletion rate almost threefold higher at 6.4 (SEM 2.4) × 10^−6^ and a gene duplication rate more than sixfold higher at 6.0 (SEM 2.6) × 10^−6^. Even after taking the average of only the nine *Con* lines with the highest CNV rates, *NiCu* lines still had a higher mean.

CNVs do not always encompass entire gene sequences, giving rise to partial gene deletions and duplications as well as complete gene CNVs. After separating these two categories, we found that rates of partial gene CNVs were generally within one order of magnitude from the rates of complete gene CNVs (Supplemental Table S1), similar to what has been found in *D. melanogaster* and *C. elegans* (Lipinski et al. 2011; Schrider et al. 2013). Overall, partial gene CNVs exhibited the same general patterns as complete gene CNVs, with significantly higher rates in *NiCu* lines (Mann-Whitney p < 0.001; Supplemental Figure S3). When only considering complete gene CNVs, *Con* lines had an average rate of 1.8 (SEM 1.0) × 10^−6^. This was not statistically different compared to both *Ni* lines at 3.9 (SEM 2.0) × 10^−6^ and *Cu* lines at 2.5 (SEM 1.3) × 10^−6^. In contrast, the *NiCu* lines had rates 3 times higher than *Con* lines (7.8 × 10^−6^; Mann-Whitney p = 0.023). These results indicate that chronic exposure to low levels of a metal mixture can substantially increase the rate at which large-scale heritable mutations arise in genomes and affect genes.

### CNV breakpoint analysis suggests a preponderance of error-prone double strand break repair

Whole genome sequencing enables nucleotide-resolution breakpoint analysis, which uses sequence information surrounding the start and end of CNVs to infer the mechanism of mutation formation such as nonallelic homologous recombination (NAHR) and nonhomologous recombination (NHR) (Lam et al. 2010). Using this approach, we found that almost every deletion (96%) was associated with the CNV formation mechanism of NHR (Supplemental Table S2). NHR consists of error-prone pathways of DNA break repair such as nonhomologous end-joining (NHEJ), and its predominance in *Daphnia* is more pronounced than what has been found in humans (Lam et al. 2010), but similar to findings in *Drosophila* (Cardoso-Moreira et al. 2012; Zichner et al. 2013). Almost half (49%) of the NHR events displayed short regions of DNA sequence homology (microhomology stretches), which is more frequent than expected based on random permutations (p = 0.002) and is a characteristic feature of NHR (Lam et al. 2010). We found that NHR events tended to have high DNA flexibility, with significantly lower helix stability (Mann-Whitney p = 0.018) and lower GC content (Mann-Whitney p = 0.016) compared to other formation mechanisms (Supplemental Figure S4). This is in line with previous findings in humans, suggesting that NHR mechanisms such as NHEJ are often associated with fragile genomic regions susceptible to double strand breaks (Lam et al. 2010). Most CNVs overlapping in multiple MA lines have different breakpoints (50-83%) suggesting independent recurrent CNVs that could represent deletion hotspots such as those previously reported in *Daphnia* (Xu et al. 2011). We found no significant differences in CNV formation mechanisms between *Con* lines and any of the metal treatments. Interestingly though, the two MA lines with the highest rates of CNVs (Con-C40 and NiCu-C376) had a CNV overlapping the *mre11* gene (Figure 3; Supplemental Table S3), a key player in DNA damage response and double strand break repair. The expression level of this gene has important consequences on the choice of DNA repair pathway and its efficiency (Rass et al. 2009). An unbalanced copy number of *mre11* could alter expression of the gene and reduce DNA repair fidelity, thereby increasing genomic CNV rates over time. We do not know, however, when the mutations to *mre11* occurred during the experiment.

### Selection against CNVs in non-MA isolates

The elevated mutation rates detected among MA lines exposed to a mixture of metals reflect heritable mutations that arise in nearly selection-free conditions, whereas purifying selection is expected to purge many new deleterious mutations. To evaluate the influence of selection on the rates and spectra of mutations in our experiment, we compared the MA lines with isolates from the population propagated under selection and seeded from the same original progenitor lineage. Given that *Con* MA lines and the non-MA population were propagated under identical environmental conditions, the underlying rate of mutation was expected to be the same, while the amount of mutations accumulated would differ due to selection. Over the same period of time as the MA experiment, the non-MA isolates accumulated 60% fewer mutations than *Con* lines, both in terms of small-scale mutations and CNVs (Supplemental Table S1). In contrast to MA lines that have each acquired mutations independently, there were three single-nucleotide mutations and two deletions detected in multiple non-MA isolates shared by descent. However, no mutations (point mutations or large-scale mutations) were shared in all isolates, suggesting that their last common ancestor is the original progenitor from the start of our experiment (Supplemental Figure S5). Accounting for the shared mutations among lineages and their genealogy, the average number of accumulated single-nucleotide mutations and CNVs in the non-MA population was three times lower than the average of MA lines. To achieve the same rate of CNVs as in *Con* MA lines, the non-MA isolates would need to have still been at generation 26 by the time *Con* lines had already reached generation 100 (Figure 5A). This very low propagation rate is however highly unlikely given that we found no difference between non-MA isolates and *Con* MA lines in either lifespan (mean of 46 days versus 43 days; t(14.6) = 1.69, p = 0.11) or age of first reproduction (mean of 11.5 days versus 11.5 days; t(14.7) = 0.18, p = 0.86). Based on mean lifespan and age of first reproduction measured in life history experiments, we calculated that the average non-MA isolate would have reached between 62 and 75 generations, with the slowest lineage at generation 30 (Supplemental Methods). Late reproduction was assigned more weight in calculating generation time, which would underestimate the number of generations if selection favored faster reproduction in the population. Nevertheless, an estimate of 62 generations still gives a mean realized rate of CNVs at least twice as low as MA lines (Table 1). Selection removing spontaneous CNVs and single nucleotide mutations is likely responsible for the lower incidence observed among non-MA isolates, as well as the lower variance observed across independent non-MA lineages compared to MA lines (Supplemental Table S1).

**Figure 5.**
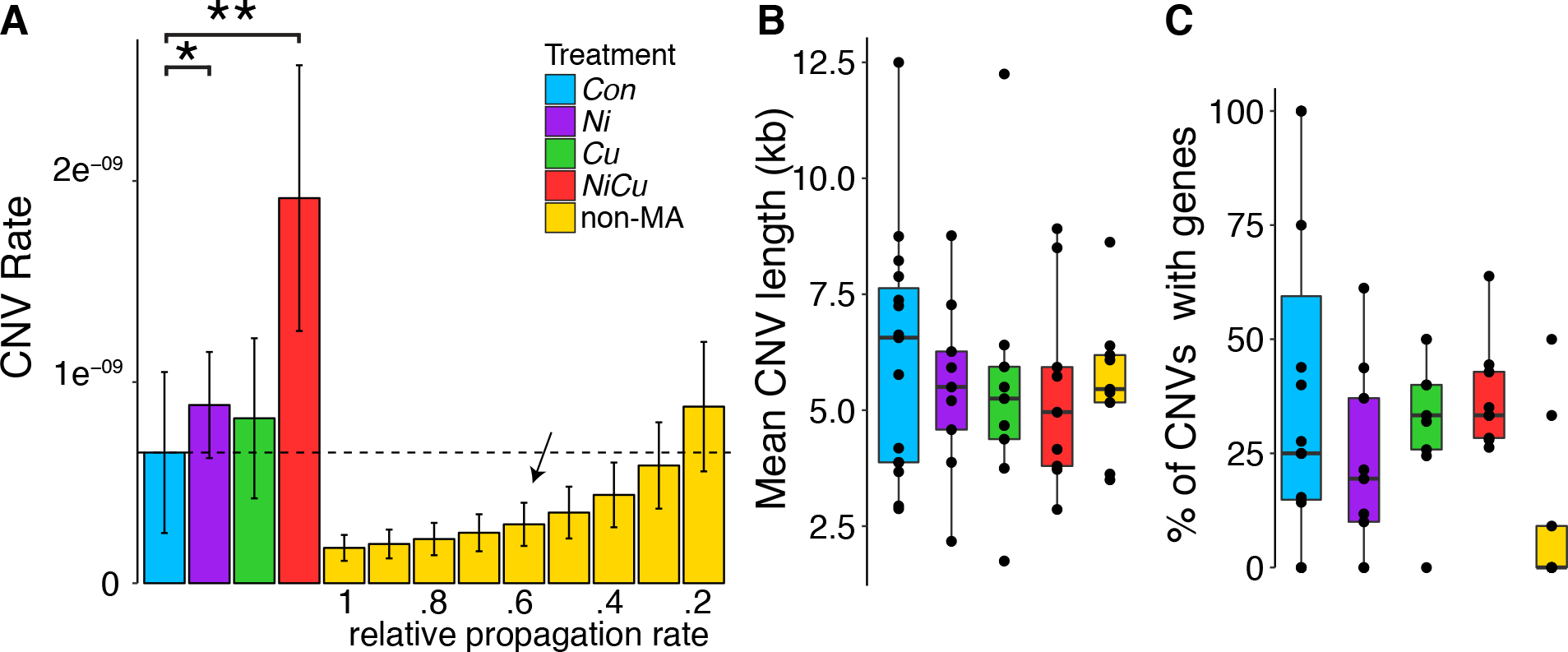
CNV rates and lengths, with and without selection. (A) Mean CNV rates (per nucleotide per generation) and standard errors among MA treatment groups showing significant differences among treatments (* for p < 0.01, ** for p < 0.005). A comparison with non-MA isolates given various relative propagation rates compared to MA lines is shown, with a dotted line indicating the relative propagation rate in non-MA genomes to reach the same mutation rates as in control MA lines. We estimated the relative propagation rate based on a life history experiment (indicated with an arrow). (B) Boxplot of CNV length distributions across treatments. (C) The percentage length of CNV regions that overlap genes, in which only a single non-MA isolate has genes deleted, and two more isolates had a gene duplication.

Given the uncertainty of the exact generation numbers in the non-MA population, characteristics of CNVs intersecting functional regions were also used to evaluate whether isolates experienced selection. We found that the proportion of CNVs that overlap single-copy genes is at least three times lower in non-MA compared to MA genomes, whereas the mean length of CNVs was not different (p > 0.05; Figures 5B and 5C). Furthermore, all but two gene CNVs were found in a single non-MA isolate (CC9) within a 100kb region deleting 7 single-copy genes (Supplemental Figure S6). Finally, there was a single partial CNV gene among non-MA genomes, while MA lines had rates of partial gene CNVs comparable to rates of complete gene CNVs (Supplemental Table 1; Supplemental Figure S3). These findings reveal a very low realized mutation rate affecting genes and the apparent efficiency of selection under constant benign conditions, suggesting that the elevated mutation rates induced by stress increase mutation burden.

## Discussion

### Comparable rates of gene deletion and duplication across organisms in benign conditions

The rates of gene deletion and duplication that we calculated using 24 MA lines of *Daphnia pulex* (3.1 × 10^−6^, or 1.8 × 10^−6^ for complete gene CNVs) are within an order of magnitude of those calculated using 8 MA lines of *S. cerevisiae* (5.5 × 10^−6^) (Lynch et al. 2008), 10 MA lines of *C. elegans* (3.4 × 10^−7^) (Lipinski et al. 2011) and 8 MA lines of *D. melanogaster* (1.1 × 10^−6^) (Schrider et al. 2013). Our rate is however much lower than a recent estimate using a similar approach and the same CNV detection software in 7 total MA lines derived from two different genetic backgrounds of *D. pulex* (5.4 × 10^−5^) (Keith et al. 2016). This difference could be partially due to the different sample sizes analyzed since we found a negative association between the number of CNVs detected and the number of genomes included in the analysis (Supplemental Figure S7). Using permutations to randomly sample the same number of genomes used in Keith *et al.* (2016), we reached a similar rate of 1.2 × 10^−5^ gene CNVs for single-copy genes among *Con* lines. However, our PCR validations confirmed the presence of false positives when analysis was performed on only a subset of our samples rather than our full dataset (Supplemental Table S4). The lower mutation rate estimate based on the full dataset is more similar to estimates from other model organisms and appears to be less susceptible to false positives in our data, although Keith *et al.* (2016) also had high validation rates for their dataset. Another difference between the two studies that could contribute to the different rates is the depth of coverage, although we found that CNV detection was not correlated with sequencing depth and that average CNV rates were unaffected after doubling the coverage of two randomly chosen MA lines. Importantly though, our study focuses on the relative mutation rates among treatments of a single genetic background, and the rate of CNVs under a metal mixture (*NiCu*) remains higher than *Con* lines regardless of the number of samples included in our analysis (either 5, 10, 20, 30 or 40 samples).

### Effects of multiple stressors on mutation rates

Multiple stressors such as metal mixtures can have different biological impacts than individual stressors, due to complex interactions (Altshuler et al. 2011; Langie et al. 2015). Here we show that the combined mixture of nickel and copper led to the highest CNV mutation rate in our experiment compared to controls, tripling rates of CNVs and quadrupling rates of gene CNVs, whereas Cu and Ni alone had a moderate to no measurable effects. This outcome is not necessarily due to the particular mixture *per se*; the *NiCu* treatment had an overall greater concentration of metals than either of the single metals alone, possibly exceeding a critical dose-response threshold. However, the similar rates of small-scale mutations across treatments suggests that DNA replication error rates and/or base and excision repair pathways were comparatively unaffected by metal stress, at least in the germ-line. This is perhaps surprising given that cellular stress can induce somatic point mutations and metals can impair excision repair pathways (Langie et al. 2015). In plants, both small-scale and large-scale mutations show higher rates under environmental stress, but different stressors have different effects: whereas high levels of salinity doubles the heritable rate of short indels and transversions (Jiang et al. 2014), various other abiotic stresses preferentially affect homologous recombination frequency compared to point mutations and microsatellite instability both in somatic cells and through transgenerational changes (Yao and Kovalchuk 2011), plausibly contributing to stress-induced CNVs (DeBolt 2010). Despite the conserved DNA repair and recombination pathways across taxa as diverged as plants and humans, species-specific duplications or deletions of genes in these pathways could contribute to differences in the repair mechanisms employed in response to DNA damage (Singh et al. 2010).

### Genetic mechanisms underlying CNVs

Given the particularly high proportion of CNVs associated with NHR across all treatments, we hypothesize that the elevated CNV rate under exposure to a metal mixture is caused by an increase in double strand breaks in the germ-line, leading to greater opportunities for recombination and DNA repair errors producing CNVs. Environmental stressors have previously been linked to increases in germ-line DNA strand breaks that are potentially caused by an increase in reactive oxygen species due to stress (Yauk et al. 2008). Additionally, metals can increase the incidence of sequence insertions at repaired double strand break sites by nonhomologous end-joining (NHEJ), proposed to be caused by interference with enzymatic processes of the proteins involved in NHEJ (Morales et al. 2016). This combination of an increase in both strand breaks and DNA repair errors could explain the higher rates of CNVs when exposed to a mixture of *Ni* and *Cu*. Although homologous recombination (HR) also leads to repair errors (Rodgers and McVey 2016), organisms that primarily repair DNA via more error-prone mechanisms might be predisposed to greater CNV rate variation and amplified effects when faced with environmental stressors. We cannot rule out the possibility that nonallelic homologous recombination (NAHR) also contributes to differences in mutation rates because our study focused on genomic regions with single-copy genes, likely underestimating the full impact of NAHR, which is an important source of recurrent CNVs occurring in segmental duplications (Gu et al. 2008).

An alternative explanation for an increase in mutation rates is that environmental stressors alter DNA repair fidelity. For example, different metals and exposure doses have been shown to differentially modulate the way cells repair double strand breaks, alternating between HR and the more error-prone NHEJ, two competing repair pathways (Morales et al. 2016). Stressful conditions in general can cause a shift to error-prone double strand break repair (Ponder et al. 2005), and lower physiological condition has been shown to lead to more mutations via changes in DNA repair pathways with different fidelity (Wang and Agrawal 2012; Sharp and Agrawal 2016). Contrary to these previous findings, we did not observe differences in the mechanism of CNV formation among treatments, potentially because CNVs under benign conditions are already associated with error-prone pathways. Instead, our results suggest that a mixture of metals induces more frequent germ-line DNA strand breakage in *Daphnia*. This finding has important evolutionary consequences particularly in taxa that propagate asexually (either cyclically or obligately) like *Daphnia*. Previous studies conducted on *Daphnia* propagated asexually under benign conditions documented high rates of loss of heterozygosity (due to gene conversion, deletion, and ameiotic recombination) that can contribute to decreasing overall fitness (Omilian et al. 2006; Xu et al. 2011; Keith et al. 2016; Flynn et al. 2017). Studying mutation rates in other organisms with different underlying genome architecture, propensity for mechanisms of DNA repair, and general levels of DNA repair fidelity would further illuminate the extent to which these genomic attributes either promote or dampen the mutagenic effects of metals in germ-lines.

## Conclusions

Despite the increasing evidence for the considerable contribution of CNVs to genome evolution and genetic diseases, few eukaryotic studies have estimated the underlying rate of large-scale duplications and deletions. Our study reports for the first time to our knowledge the variation in genome-wide rates of CNVs under different environmental conditions, as well as under different selection regimes using the same genetic background. Our results illustrate that exposure to low but ecologically relevant levels of metal mixtures can accelerate rates of large-scale deletions and duplications, likely increasing mutational load and triggering deleterious phenotypic effects. Sequence analyses at CNV breakpoints suggest that low levels of metals induce germ-line DNA strand breakage rather than modify DNA repair pathways used for resolving double strand breaks. Chronic exposure to low environmental stress might therefore have profound consequences on the frequency and type of variation generated in the genome, including duplicated and deleted genes, ultimately influencing the mutation burden and evolutionary trajectory of natural populations.

## Methods

### Daphnia mutation accumulation experiment

To assess mutation rate variation under different environmental conditions, we conducted a mutation accumulation (MA) experiment using a total of 51 independent lines of *Daphnia pulex* over an average of 103 generations (Figure 1). Twenty-four replicate MA lines were propagated in benign soft-water media as described in Flynn *et al.* (2017), herein labeled as MA controls (*Con*). In addition, nine nickel-exposed MA lines (*Ni*) were maintained in 80 μg/L of nickel, nine copper-exposed MA lines (*Cu*) were maintained in 40 μg/L of copper, and nine MA lines were maintained in a mixture of nickel and copper (*NiCu;* 80 μg/L of nickel + 40 μg/L of copper). These sub-lethal concentrations of metals did not elicit a measurable differences in mortality, average brood size, or time to first clutch, and are comparable to *Daphnia* habitats that experienced decades of contamination of copper and nickel in the Sudbury Canada area (Yan et al. 2016). Each MA line was propagated using single progeny descendants every generation, and all were seeded with a single *Daphnia* obligate parthenogenetic progenitor. The ancestral progenitor for all MA lines was collected from Canard Pond (Lat. 42°12”, Long. -82°98”) in Windsor Ontario, Canada. All MA lines were maintained at 18°C with a humidity of 70% and a 12 hour light/dark cycle. MA lines were fed *ad libitum* with a mixture of algae (*Ankistrodesmus sp., Scenedesmus sp.* and *Pseudokirchneriella sp).* Backups for MA lines were maintained in case of mortality or sterility of the focal individual, and were used in ~6% of transfers with an average of once every 16 generations per line. Although this introduces some level of selection against lethal and sterility-causing mutations that could lead to underestimating mutation rates, the frequency of backup lines used across treatments was not significantly different.

### Daphnia population under selection

To evaluate the effect of selection on mutation rates, a large non-MA population seeded from the same ancestral progenitor as the MA lines was allowed to propagate without induced population bottlenecks for the duration of the MA experiment. Thus, whereas the MA lines experienced minimal selection, the non-MA population evolved with selection. The non-MA population was maintained in a 15 L tank under the same conditions as the *Con* MA lines with identical media, temperature and lighting conditions. Feeding was performed twice a week using the same mixture of algae as the MA lines. Six isolates were randomly chosen for sequencing when the *Con* MA lines had reached an average of 101 generations (1,368 days of propagation), and three additional isolates were sequenced after an average of 136 generations (1,642 days of propagation; see Supplemental Methods). The census size of the population at the earlier time point was estimated to be between 100 and 250 (Flynn et al. 2017). Although natural population size fluctuations probably occurred, the lack of fixed mutations (i.e. shared across all non-MA isolates versus the ancestor) and the few shared mutations observed provides little evidence for severe population bottlenecks (Supplemental Figure S5). Future studies involving highly replicated non-MA populations would be needed to assess the extent of stochastic allele frequency variation among different populations.

### Sample processing and sequencing

Tissue collection, library preparation, and sequencing followed the approach described in Flynn *et al.* (2017). Tissue was collected from 3-5 clonal individuals per line raised in a sterile medium. During 48 hours prior to isolating DNA, animals were fed sterile Sephadex beads 10 times a day to eliminate food content from the gut, while being treated with antibiotics to reduce microbial contamination. DNA was extracted following the cetyltrimethylammonium bromide method (Doyle and Doyle 1987). DNA samples were quantified with PicoGreen Quant-IT and were diluted to 2.5 ng/μL. We adopted a library preparation protocol derived from the standard Illumina Nextera approach that was optimized to reduce the use of reagents (Baym et al. 2015). Samples were dual indexed (one index at the 3’ end and another index at the 5’ end) such that each sample had a unique index combination per sequencing lane. Libraries were cleaned and short products removed with Beckman Coulter AMPure XP beads. Libraries were then normalized, pooled into three groups, and run on a total of five lanes of Illumina HiSeq 100bp and 150bp paired end reads at Genome Quebec. Adapter sequences were removed and overlapping sequences merged from fastq files using SeqPrep (https://github.com/jstjohn/SeqPrep). For each of the sequencing lanes, reads were mapped against the *Daphnia pulex* reference genome (Colbourne et al. 2011) using the short read alignment tool BWA v0.7.10 (Li and Durbin 2009). After alignment, reads were cleaned and sorted, and duplicates were removed with Picard tools v1.123 (http://broadinstitute.github.io/picard). Resulting BAM files were used for estimating depth of coverage, achieving an average of 13x coverage. Two randomly selected MA lines (C01 and C35) were intentionally sequenced to a higher depth to test the effect of doubling the sequence coverage on mutation rates. All analyses were carried out twice, once before the increase in coverage of the two samples, and once after the increase in coverage. This increase in coverage did not affect the recovery of CNVs, nor the estimated mutation rate of single nucleotides (Flynn et al. 2017).

### Small-scale variant calling

Single nucleotide mutations and INDELs were called using GATK v.3.3.0 (McKenna et al. 2010), first using HaplotypeCaller to assign putative genotypes for each individual separately, followed by GenotypeGVCFs to refine variant calling over all samples simultaneously. Variants were filtered using GATK based on various quality and alignment metrics including variant quality, mapping quality, and strand bias (QD<2, QUAL<50, FS>60, MQ<40, MQRankSum<-12.5, ReadPosRankSum<-8 for single nucleotide mutations, and QD<2, QUAL<50, FS>200, ReadPosRankSum<-20 for INDELs). To further prevent false positive variant calls from the sequencing data, we excluded non-nuclear sites, repeat masked regions, sites without read coverage from each sample, and regions with overall depth lower than expected (average 6x) or greater than twice the expected coverage (average 26x). These filtering steps were informed by both follow-up inspection of mapped reads in a genome browser and Sanger sequencing of single nucleotide mutations and INDELs called at various filtering stages and with different read depths as described in Flynn *et al.* (2017). We retained ~25% of the reference genome as callable sites for identifying single nucleotide mutations and INDELs. As expected, all MA lines had unique mutation profiles despite allowing shared mutations among lines. We did not identify shared single nucleotide mutations across MA lines or any signature of potential contamination across lines propagated in isolation. The raw sequence data can be found in SRA (PRJNA341529).

### Large deletions and duplications

Four different CNV detection programs were initially run for determining putative deletions and duplications utilizing read depth, split-read and/or paired-end approaches. Read depth analysis was performed using CNVnator v0.3 (Abyzov et al. 2011) with a bin size of 500bp to uncover putative deletions and duplications for each sample compared to the reference genome. Another read depth approach called CNV-seq v0.2-8 (Xie and Tammi 2009) was used that compares pairwise samples. CNV-seq was run using a sliding 250bp window on every pairwise comparison between MA lines (i.e. all pairwise combinations among the 51 *Con, Ni, Cu,* and *NiCu* samples), and between each non-MA isolate and every MA line (but not non-MA isolates with one another since they can share CNVs by descent). CNVs were called if four consecutive windows had a log_2_ depth of coverage difference above 0.44 or below 0.6, which requires a coverage ratio >1.36 or <0.66 respectively. CNVs detected in every pairwise comparison were identified for each sample, followed by the merging of CNVs within 10kb of one another to represent a single CNV, to overcome the majority of assembly gaps and repetitive regions (Keith et al. 2016). Paired-end read mapping and soft-clipped split-reads were also used to infer structural variants using SoftSV v1.4 (Bartenhagen and Dugas 2016). Because paired reads of short fragments overlap one another, inhibiting the ability to detect CNVs, SoftSV was also analyzed after trimming all reads to 50bp. Trimming the ends of paired reads can theoretically permit independent mapping of each paired read by removing overlapping sequences, thereby improving chances of detecting CNVs. In addition to the three tools mentioned above, we used a simple in-house read depth approach to estimate CNVs among genes in individual lines as follows. For each gene from each sample, read depth was standardized by the total read depth of the respective sample to enable comparisons across lines, and read depth was centered to 2 to approximate diploid copy numbers. We compared all MA lines with one another and with non-MA isolates using the deviation of normalized read depth among samples to identify candidate gene duplications and gene deletions. At a diploid locus, we would expect mutants with a deletion to have at least half as much coverage as non-mutant lines, and mutants with a duplication to have at least twice as much coverage as non-mutant lines. Due to variability in read depth coverage, we used slightly less stringent thresholds while still requiring mutants to be outliers based on 1.5x interquartile ranges. Genes were considered as deleted if the MA line with the lowest standardized read depth was < 0.66x compared to all other MA lines, while being an outlier with at least 0.5 fewer absolute copies. Genes were considered as duplicated if the MA line with the highest standardized read depth was > 1.4x compared to all other MA lines, while being an outlier with at least 0.5 more absolute copies. Based on the overlaps of CNVs detected from all four methods, CNV-seq had an overwhelmingly higher proportion of gene CNVs overlapping our read depth method (up to 10 fold more than both CNVnator and SoftSV), and also shared the highest proportion of pairwise concordant CNVs among the 3 implemented tools. Combined with the fact that CNV-seq was also used in a recent analysis among other *Daphnia* MA lines and had high validation rates (Keith et al. 2016), we decided to solely rely on the results of CNV-seq. To evaluate the effects of sample size on CNV detection, we repeated our CNV analyses and rate estimates using random sampling of 5, 10, 20, 30 and 40 genomes. Whereas absolute rate estimates differed, the relative rates between treatments were not affected.

To identify CNVs shared by descent as well as mutation hotspots, we allowed overlapping (shared) CNVs among samples (e.g. two samples with deletions versus all other samples but not between one another). However, we did not allow shared mutations to occur in more than 50% of lines, which could be due to differences between the ancestor and the reference genome. Overlapping CNVs in non-MA isolates were interpreted as shared by descent, and shared CNVs among MA lines were considered as recurrent CNVs (potentially hotspots). CNVs with an average depth of coverage below 6x were removed. Protein-coding genes that intersected with remaining duplications or deletions (with a minimum 5% of their length) were considered as putative gene CNVs (>95% length overlap were considered as “complete” gene CNVs as opposed to partial gene CNVs).

### Mutation validations

Sanger sequencing of randomly selected single nucleotide mutations and INDELs confirmed 21 out of 25 mutations as described in Flynn *et al.* (2017). Long-range PCR amplification of CNVs was performed to validate the presence or absence of large-scale mutations in the putative mutant sample and two other independent MA lines. Primer pairs were designed based on the ancestral progenitor’s sequence around inferred breakpoints from randomly selected CNV loci, in addition to one CNV overlapping the *mre11* gene, two CNVs found in multiple samples, and five CNVs that were excluded after filtering (Supplemental Table S4). Our PCR approach successfully verified 12 out of 14 CNV tests, and confirmed all (4 out of 4) putative CNVs that were called with fewer samples (but not detected after increasing the number of sample comparisons) were false positives.

### CNV rate calculation

Duplications and deletions were evaluated using only the scaffolds that contained one of the 10,673 “single-copy” protein-coding genes in *Daphnia* to reduce the impact of mis-mapping against the highly duplicated reference genome (Colbourne et al. 2011; Keith et al. 2016). Single-copy genes were determined as genes without any duplicates in the *Daphnia* reference genome by identifying paralogs using EnsemblMetazoa v30. The number of sites kept for analysis and used to calculate mutation rates was 113,196,346bp (57% of the reference genome), with 8,699 single-copy protein-coding genes found on 1,313 scaffolds. CNVs that had an average coverage below 6x across all samples were removed. The duplication and deletion rates per genome were estimated using the formula *μ* = *n / T*, where *n* equals the number of duplication or deletion events and *T* is the number of generations that a sample was propagated. Since all samples are compared over the same genomic regions, these rates can be used to compare treatments. CNV rates per genome per nucleotide were compared across studies and were calculated using *μ* = *n* / (2 × *L* × *T*), in which *L* is the total number of loci (nucleotides) analyzed. For mutation rates of gene duplications and deletions, *n* was the number of gene CNVs and *L* was the number of single-copy protein-coding genes analyzed as mentioned above.

The nine non-MA isolates were sampled at two time points: six when MA *Con* lines reached an average of 101 generations (1,368 days), and three more when MA *Con* lines reached an average of 136 generations (1,642 days). Due to potentially overlapping generations, the non-MA population likely achieved lower mean generations than MA-lines. To estimate the average number of generations, ten non-MA isolates and ten *Con* MA lines with seven replicate offspring from each focal mother were used in a life history experiment (Supplemental Methods). The average generation time of the population was estimated based on the mean age at first reproduction and longevity, and weighted by average clutch sizes. To compare realized rates of CNVs in the non-MA population that has likely faced greater selective pressures than the MA lines, we calculated the realized mutation rates taking into account the genealogy and shared mutations among lineages (Supplemental Methods). Moreover, we used a range of generations in the denominator to represent propagation rates up to five times slower relative to MA lines (i.e. relative propagation rates from 0.2 to 1), encompassing the average and lower bound generation estimates.

### Breakpoint detection and deletion formation mechanisms

Breakpoint analysis of CNVs was performed to infer the molecular mechanism of deletion formation. This approach compares the nucleotide sequences surrounding the ends (breakpoints) of CNVs with expected genomic signatures of different CNV formation mechanisms, including the identification of transposable elements, repeats, low-complexity DNA motifs, and sequence identity across breakpoint junctions (Lam et al. 2010). For each deletion, reads that mapped around putative breakpoint boundaries were assembled using TigraSV v0.4 (Chen et al. 2014). Assembled contigs were then re-aligned to the reference to define breakpoints using AGEv0.4 (Abyzov and Gerstein 2011). Filtering was performed to assign breakpoints with high confidence based on the comparison between the sequence alignment and the predicted deletion region; contig alignments were required to have at least 95% sequence identity including flanking regions, as well being within 4kb of the estimated breakpoint ranges from CNV-seq and overlapping at least 50% of the estimated range. When multiple alternative breakpoints were found, we selected the ones closest to the estimated range. Breakpoints were estimated for each CNV, and deletion formation mechanisms (such as NHR) were inferred for 43% of deletions using BreakSeq v1.3 (Lam et al. 2010) as well as the DNA flexibility, DNA helix stability, and GC content at breakpoints.

### Data access

The genome sequence data for all samples have been deposited in the Sequence Read Archive (SRA) under accession PRJNA341529.

## Acknowledgments

We thank all the students that contributed to maintaining the MA lines over the past five years. Thanks to T. Crease, D. Denver, B. Fryer, D. Haffner, R. Vergilino, K. Millette, N. Yan, J. Witt, G. Zhang and M. Dutton for helpful suggestions during the project.

## Author contributions

FJJC and MEC conceived and designed the project. JMF and JKB prepared the samples for sequencing. JMF and FJJC designed the analytical approach and performed the PCR validations. JKB performed the life history experiment. FJJC analyzed the data and wrote the paper. All authors contributed to and approved the final manuscript. This project was supported by an NSERC CREATE fellowship to FJJC, an NSERC scholarship to JMF, and an NSERC CREATE training program on Aquatic Ecosystem Health, an NSERC Discovery Grant, and Canada Research Chair to MEC.

